# Chromosome-level genome assembly of the African pike, *Hepsetus odoe*

**DOI:** 10.1101/2020.05.13.094987

**Authors:** Xiao Du, Xiaoning Hong, Guangyi Fan, Xiaoyun Huang, Shuai Sun, Ouyang Bingjie, He Zhang, Mengqi Zhang, Shanshan Liu, Xin Liu, Wenwei Zhang

## Abstract

The order Characiformes is one of the largest components of the freshwater teleost fauna inhabiting exclusively in South America and Africa with great ecological and economical significance. Yet, quite limited genomic resources are available to study this group and their transatlantic vicariance. In this study we present a chromosome-level genome assembly of the African pike (*Hepsetus odoe*), a representative member of the African Characiformes. To this end, we generated 119, 11, and 67 Gb reads using the single tube long fragment read (stLFR), Oxford Nanopore, and Hi-C sequencing technologies, respectively. We obtained an 862.1 Mb genome assembly with the contig and scaffold N50 of 347.4 kb and 25.8 Mb, respectively. Hi-C sequencing produced 29 chromosomes with 742.5 Mb, representing 86.1% of the genome. 24,314 protein-coding genes were predicted and 23,999 (98.7%) genes were functionally annotated. The chromosomal-scale genome assembly will be useful for functional and evolutionary studies of the African pike and promote the study of Characiformes speciation and evolution.

## Background & Summary

The order Characiformes is one of the largest components of the freshwater fish fauna worldwide, comprising about 2,000 ecologically and morphologically diverse fish living in rivers and lakes exclusively in Africa and South America. Characiformes are ecologically important by playing crucial roles in energy flux and material cycling in river systems^1^. Moreover, they have great significance for local economy because of diet component of livestock and humans^2^. As Characiformes are exclusively freshwater fishes, their transatlantic distribution was proposed ascribed to the split of South America and Africa in the Early Cretaceous fragmentation of western Gondwana^3^. This distribution across the Atlantic Ocean displays asymmetry in the number of species, with *circa* 220 reported species in Africa and over 1,700 species in South America. High fragmentation has been reported in the South American species, compared to the lower fragmentation and variability in African ones^4,5^. However, quite limited genomic information has been available to study the Characiformes vicariance. Despite the large amount and high diversity of species, presently only three genomes from two families *Characidae* (*Astyanax mexicanus*) and *Serrasalmidae* (*Pygocentrus nattereri, Colossoma macropomum*) have been released in Characiformes, which all belong to the South American lineages. No genomes of African Characiformes have been reported. Genomic studies of African Characiformes would highly promote the understanding of Characiformes evolution and speciation during the continent fragmentation.

African pike, *Hepsetus odoe*, is a representative African Characiformes that belongs to the family *Hepsetidae*. It is a torpedo-shaped predatory and piscivorous species distributed in the freshwater basins in central and western Africa^6^, bearing a striking resemblance to the European pike. One of the most striking features is their dentition with the lower jaw filled with two rows of sharp point teeth while the upper with only one raw. Due to roles in freshwater food chain and diet component of livestock and humans, *H*.*odoe* are biologically and economically important^2^. *H*.*odoe* was reported the only species in the *Hepsetidae* family, until recently five additional species were described by recent studies^6,7^. Despite of the high economic and evolutionary importance, no genome data are available for this group.

A high-quality genome assembly of the African pike will highly facilitate the study of its functional and evolutional genomics, which also will promote the understanding of other African Characiformes along with their divergence from the South American Characiformes. Therefore, in this study we report a chromosome-scale genome assembly of *H*.*odoe* using single tube long fragment read (stLFR)^8^, Oxford Nanopore, and Hi-C technologies (Additional file 1: Fig. S1). We obtained a genome assembly of 862.1 Mb with the contig and scaffold N50 of 347.4 kb and 25.8 Mb, respectively. With chromosome-level scaffolding, 29 scaffolds were constructed corresponding to 29 chromosomes with a total length of 742.5 Mb, representing 86.1% of all genome sequences. 24,314 protein-coding genes were predicted in the assembly, and 98.7% of them were functionally annotated. The chromosomal-level genome assembled here will be useful for functional and evolutionary research of the African pike. It is the first genome assembly in the African Characiformes and will promote the understanding of Characiformes speciation and evolution.

## Methods

### Sampling and sequencing

Long genomic DNA (gDNA) from muscle tissue of a male African pike was isolated using a conventional approach for sufficient DNA quality^9^. DNA integrity was checked using agarose gel electrophoresis. The sequencing libraries were constructed via stLFR technology according to the standard protocol via the MGIEasy stLFR library preparation kit (PN:1000005622)^10^ and were sequenced on BGISEQ-500 platform. To overcome the gaps (long ambiguous sequences) induced by repeats, library preparation and sequencing were performed on the MinION nanopore sequencer (Oxford Nanopore Technologies, Oxford, UK) for generating long reads, following the base protocols of Oxford Nanopore. To get a high-resolution genome contact map, we used *in situ* Hi-C according to the protocol of previous study with some modifications^11^. The restriction endonuclease MboI was used to digest DNA, followed by biotinylated residue labeling. The Hi-C library was sequenced on BGISEQ-500 platform with 100 bp pair-end sequencing.

### Ethics statement

The adult male African pike was purchased from the fish and aquarium market in Guangzhou, Guangdong Province, China in May 2018. The experimental procedures followed the guidelines approved by the institutional review board on bioethics and biosafety of BGI (IRB-BGI). The experiment was authorized by the IRB-BGI (under NO. FT17007). The review procedures in IRB-BGI meet good clinical practice (GCP) principles.

### *De novo* assembly, and chromosome construction

The k-mer frequency distribution analysis^12^ was used to estimate the African pike genome size. According to the 17-mer analysis, the genome size of African pike was estimated to be 995 Mb (Table 2; Additional file 1: Fig. S2).

**Table 1.** Sequencing results for African pike genome assembly.

| Libraries | Raw data |  | High-quality data |  |
| --- | --- | --- | --- | --- |
|  | Total bases | Sequencing depth | Total bases | Sequencing depth |
|  | (Gb) | (×) | (Gb) | (×) |
| stLFR | 118.6 | 141.2 | 60.38 | 71.88 |
| Nanopore | NA | NA | 11.02 | 13.12 |
| Hi-C | 66.86 | 77.47 | 65.82 | 76.25 |

**Table 2.**
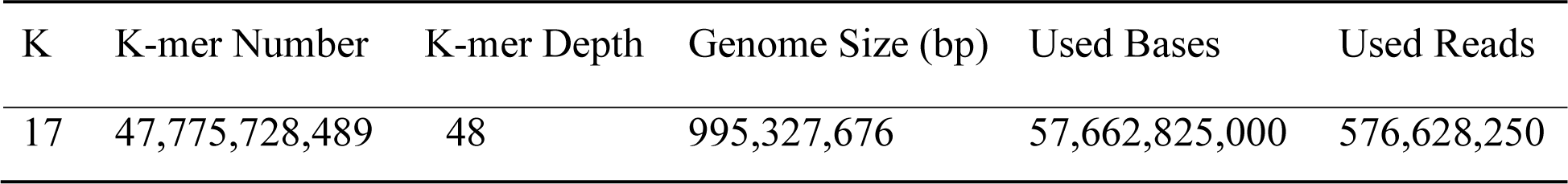
K-mer analysis for African pike genome.

| K | K-mer Number | K-mer Depth | Genome Size (bp) | Used Bases | Used Reads |
| --- | --- | --- | --- | --- | --- |
| 17 | 47,775,728,489 | 48 | 995,327,676 | 57,662,825,000 | 576,628,250 |

We obtained 118.6 Gb (∼141X; Table 1) raw sequencing reads from stLFR. We used SOAPfilter v.2.2, a package in SOAPdenovo2^13^ to filter reads with low quality reads (> 40% low-quality bases, Q <7), PCR duplication, or adapter contamination. After filtering, 60.4 Gb (∼72X; Table 1) clean reads were obtained for genome assembly. Supernova assembler v2.0.1 (10X Genomics, Pleasanton, CA) was used to build contigs and scaffolds, and gaps were closed by GapCloser (v1.2)^13^. With stLFR data, the generated African pike assembly was 859.2 Mb. The contig and scaffold N50 were 43.9 kb and 5.1 Mb, respectively (Table 3). On basis of that, we generated a total of 11.0 Gb (∼13X; Table 1) long reads on the MinION nanopore sequencer to further fill the gaps using TGSGapFiller^14^ with default parameters. After gap filling, the contig length was highly elevated with contig N50 of 352.1 kb (Table 3).

**Table 3.**
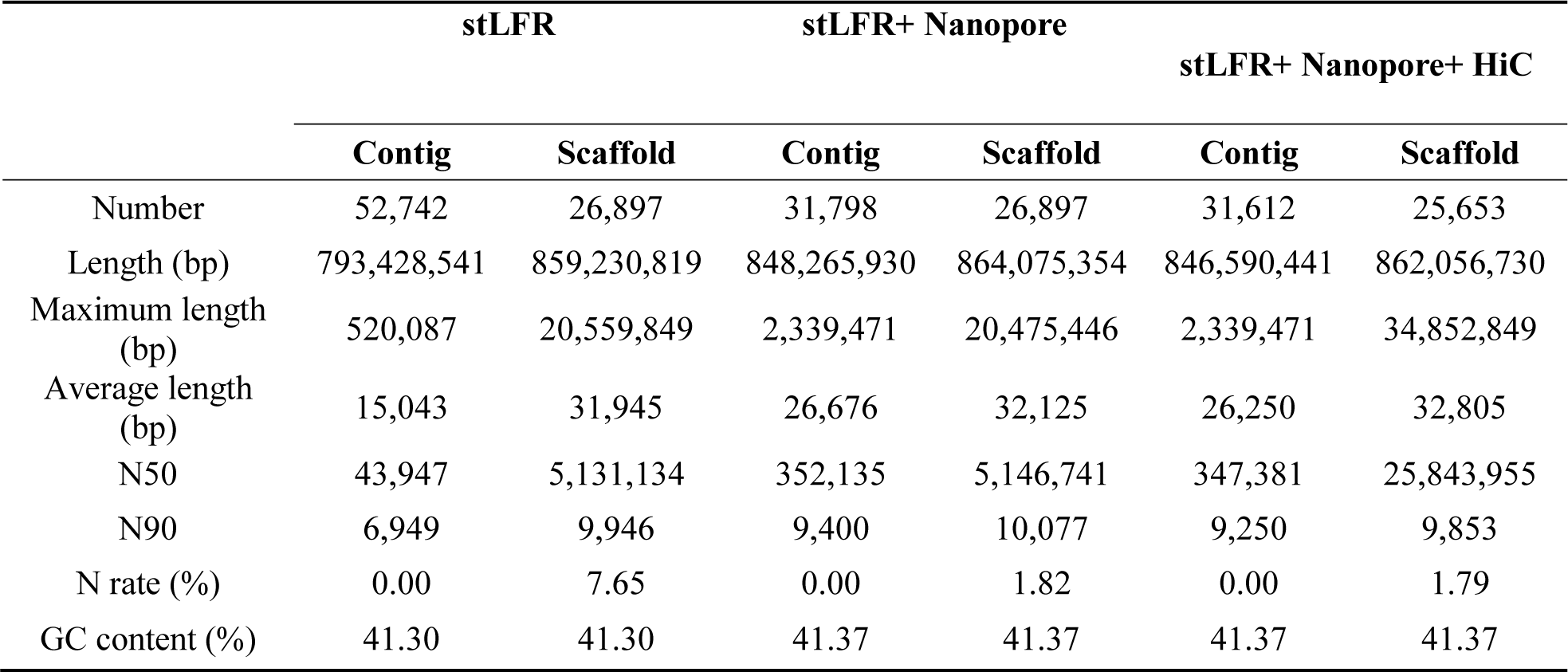
Assembly statistics for the African pike.

|  | stLFR |  | stLFR+ Nanopore |  | stLFR+ Nanopore+ HiC |  |
| --- | --- | --- | --- | --- | --- | --- |
|  | Contig | Scaffold | Contig | Scaffold | Contig | Scaffold |
| Number | 52,742 | 26,897 | 31,798 | 26,897 | 31,612 | 25,653 |
| Length (bp) | 793,428,541 | 859,230,819 | 848,265,930 | 864,075,354 | 846,590,441 | 862,056,730 |
| Maximum length (bp) | 520,087 | 20,559,849 | 2,339,471 | 20,475,446 | 2,339,471 | 34,852,849 |
| Average length (bp) | 15,043 | 31,945 | 26,676 | 32,125 | 26,250 | 32,805 |
| N50 | 43,947 | 5,131,134 | 352,135 | 5,146,741 | 347,381 | 25,843,955 |
| N90 | 6,949 | 9,946 | 9,400 | 10,077 | 9,250 | 9,853 |
| N rate (%) | 0.00 | 7.65 | 0.00 | 1.82 | 0.00 | 1.79 |
| GC content (%) | 41.30 | 41.30 | 41.37 | 41.37 | 41.37 | 41.37 |

Reads from Hi-C library^15^ were used to generate a chromosomal-level genome assembly. First, we obtained 65.8 Gb (∼76X, Table 1) clean sequencing data from the Hi-C library by removing reads containing more than 1% unidentified (N) bases and low-quality bases (quality value < 10) using SOAPnuke (v1.5.4)^16^ with parameters “-l 10 -q 0.1 -n 0.01 -Q 2”. Next, we used HiC-Pro pipeline (v2.8.0)^17^ for quality control to generate valid reads. Of all 658,260,000 raw pair-end reads, there were 22.78% valid (149,912,370) paired Hi-C reads suitable for following analysis. We used Juicer (v.1.5)^18^, an open-source and fully-automated pipeline for pretreatment of Hi-C datasets, for analyzing valid Hi-C datasets and producing the alignment result. Lastly, we applied 3D-DNA workflow (3D *de novo* assembly, v.170123)^19^ to create the ordered-and-oriented genome sequences in chromosome level with the main parameter “-m haploid -s 4 -c 29”. We assembled 29 chromosomes of *H. odoe* ranging from 8.03 Mb to 34.85 Mb with the total length of 742.5 Mb (Table 4; Fig. 1), which possessed 86.1% of all genome sequences. The final African pike genome assembly spanned 862.1 Mb and 29 chromosomes, accounting for 86.6 % of the estimated genome size, with contig and scaffold N50 of 347.4 kb and 25.8 Mb, respectively. The constructed 29 chromosomes agreed with the previous karyotype analysis of *H. odoe*^6^.

**Table 4.** Summary of the assembled 29 chromosomes in the African pike.

| Chromosome | Number of contigs | Length of contigs(bp) |
| --- | --- | --- |
| 1 | 174 | 34,852,849 |
| 2 | 155 | 33,370,094 |
| 3 | 191 | 32,320,846 |
| 4 | 163 | 31,998,173 |
| 5 | 169 | 31,241,732 |
| 6 | 150 | 30,610,121 |
| 7 | 231 | 30,524,172 |
| 8 | 175 | 29,799,086 |
| 9 | 132 | 29,196,192 |
| 10 | 157 | 29,175,684 |
| 11 | 145 | 27,911,487 |
| 12 | 133 | 27,279,904 |
| 13 | 104 | 26,912,730 |
| 14 | 144 | 26,647,357 |
| 15 | 118 | 25,843,488 |
| 16 | 137 | 25,843,955 |
| 17 | 130 | 25,569,178 |
| 18 | 131 | 25,245,420 |
| 19 | 122 | 24,922,465 |
| 20 | 153 | 24,594,709 |
| 21 | 216 | 23,062,401 |
| 22 | 116 | 22,653,434 |
| 23 | 166 | 22,465,777 |
| 24 | 82 | 21,478,618 |
| 25 | 168 | 21,151,346 |
| 26 | 121 | 19,748,208 |
| 27 | 135 | 18,283,571 |
| 28 | 48 | 11,790,282 |
| 29 | 76 | 8,027,360 |
| Total | 4,142 | 742,520,639 (86.1%) |

**Fig. 1.**
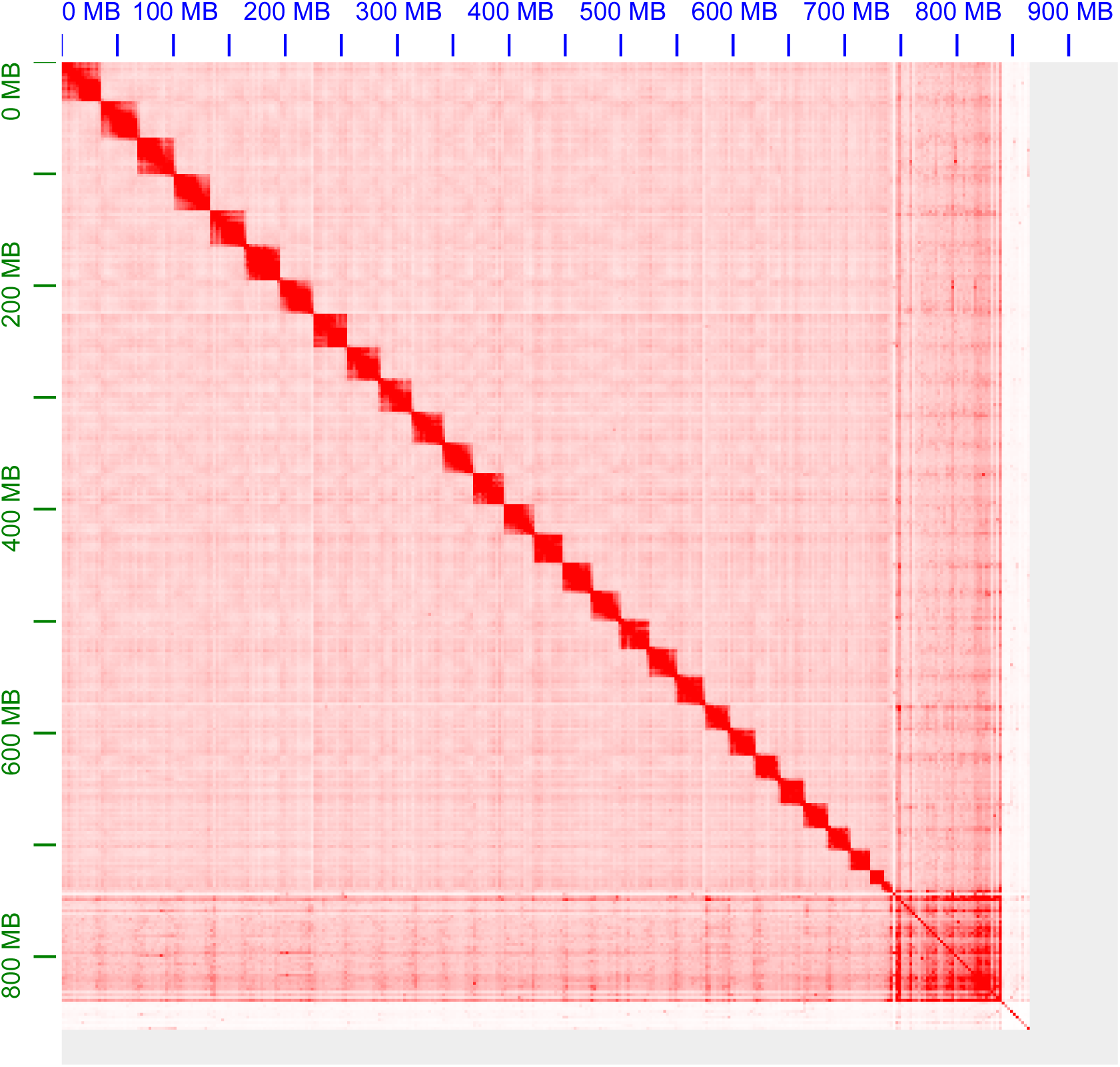
Hi-C chromosome heat map of African pike genome. Each block represents a Hi-C contact between two genomic loci within a 100kb bin. Darker color represents higher contact intensity.

### Gene prediction and functional annotation

To facilitate gene prediction in the genome, repetitive elements were identified first. Two methods (*de novo* and homology-based predictions) were performed in the repeat annotation of the African pike genome. In the *de novo* method, a *de novo* library was built via running RepeatModeler (v1.0.8)^20^ and LTR-FINDER (v1.0.6)^21^, and the predicted model was applied to identify interspersed repetitive elements by RepeatMasker (v4.0.5). In the homology-based prediction, detection of interspersed repeats was realized by aligning the genome against the Repbase database^22^ at DNA and protein levels using RepeatMasker and RepeatProteinMask (v4.0.5)^23^. Tandem repeats were predicted by TRF (v4.07). By integrating results of above approaches, 317.3 Mb repetitive sequences were predicted, representing 36.7% of the genome assembly (Table 5). Finally, 284.1 Mb TEs were identified, accounting for 32.9% of the genome assembly. The repetitive element annotations were summarized in Table 6. Those repetitive sequences were masked to reduce the interference for the following gene predictions.

**Table 5.** Prediction of repetitive sequences in African pike.

| Type | Repeat Size (bp) | % of genome |
| --- | --- | --- |
| TRF | 42,227,324 | 4.89 |
| RepeatMasker | 140,715,741 | 16.30 |
| RepeatProteinMask | 36,712,306 | 4.25 |
| *De novo* | 292,061,790 | 33.81 |
| Total | 317,311,789 | 36.73 |

**Table 6.** Repeat annotation of the African pike assembly.

| Type | RepBase TEs |  | TE Proteins |  | *De novo* |  | Combined TEs |  |
| --- | --- | --- | --- | --- | --- | --- | --- | --- |
|  | Length (bp) | % in genome | Length (bp) | % in genome | Length (bp) | % in genome | Length (bp) | % in genome |
| DNA | 64,714,586 | 7.49 | 2,115,720 | 0.25 | 106,985,346 | 12.39 | 140,841,144 | 16.30 |
| LINE | 49,378,300 | 5.72 | 31,164,294 | 3.61 | 134,752,407 | 15.6 | 153,325,343 | 17.75 |
| SINE | 25,662,188 | 2.80 | 0.00 | 0.00 | 10,933,947 | 1.27 | 34,624,631 | 4.00 |
| LTR | 15,943,766 | 1.85 | 3,844,230 | 0.45 | 67,207,893 | 7.78 | 77,013,770 | 8.92 |
| Other | 17,549 | 0.002 | 0.00 | 0.00 | 0.00 | 0.00 | 17,549 | 0.002 |
| Unknown | 0.00 | 0.00 | 0.00 | 0.00 | 3,180,101 | 0.37 | 3,180,101 | 0.37 |
| Total | 140,715,741 | 16.30 | 36,712,306 | 4.25 | 268,746,546 | 31.12 | 284,106,647 | 32.89 |

Next, we conducted structural and functional annotation for the assembled genome. For structural annotation, both homology-based and *de novo* prediction approaches were applied. In *de novo* prediction, AUGUSTUS (v3.1)^24^ and GENSCAN (v2009)^25^ were utilized to predict the gene model with zebrafish data as a training set, and 23,163 and 29,084 protein-coding genes were predicted, respectively (Table 7). The homology-based prediction of genome assembly was realized by referring to the NCBI protein repertoires of six homologous species including Mexican tetra (*Astyanax mexicanus*), red-bellied piranha (*Pygocentrus nattereri*), channel catfish (*Ictalurus punctatus*), common carp (*Cyprinus carpio*), iridescent shark *(Pangasianodon hypophthalmus)*,and zebrafish (*Danio rerio*). After mapping the protein sequences to the repeat-masked African pike genome using BLAST^26^ (*E*-value cutoff of 1xE^-5^), GeneWise (v2.4.1)^27^ was used to predict gene models by aligning homologous genome sequences against the matched proteins. Lastly, we preformed GLEAN to integrate all above gene models and obtained a non-redundant gene set consisting of 24,314 protein-coding genes (Table 7). There were 9.77 exons per gene and the average length of coding sequences (CDS) was 1,712 bp (Table 7). Gene function was annotated with TrEMBL^28^, Swissprot^28^, InterPro^29^, Gene Ontology^30^, and Kyoto Encyclopedia of Genes and Genomes (KEGG)^31^ databases. Ultimately, 23,999 genes (98.7% of the total) in African pike were functionally annotated (Table 8).

**Table 7.** Statistics of gene annotations in African pike assembly.

| Gene set |  | Number | Average transcript length (bp) | Average CDS length (bp) | Average exon per gene | Average exon length (bp) | Average intron length (bp) |
| --- | --- | --- | --- | --- | --- | --- | --- |
| *De novo* | Augustus | 23,163 | 14,343 | 1,392 | 7.95 | 175 | 1,862 |
|  | Genscan | 29,084 | 18,389 | 1,461 | 8.06 | 181 | 2,398 |
| Homolog | *I.punctatus* | 45,520 | 27,590 | 2,021 | 11.57 | 175 | 2,418 |
|  | *D.rerio* | 26,485 | 15,268 | 25,114 | 8.68 | 185 | 3,060 |
|  | *P.nattereri* | 45,066 | 29,558 | 1,968 | 11.22 | 175 | 2,699 |
|  | *P.hyp* | 37,334 | 29,021 | 2,084 | 11.95 | 175 | 2,461 |
|  | *A.mexicanus* | 41,543 | 32,069 | 1,991 | 11.54 | 173 | 2,855 |
|  | *C.carpio* | 55,544 | 13,199 | 1,176 | 6.72 | 175 | 2,100 |
| Combined | GLEAN | 24,314 | 15,835 | 1,712 | 9.77 | 175 | 1,610 |

**Table 8.** Functional annotations of predicted genes in African pike assembly.

|  | Database | Number | Percentage (%) |
| --- | --- | --- | --- |
| Total |  | 24,314 | 100 |
|  | InterPro | 23,989 | 96.35 |
|  | GO | 17,227 | 70.85 |
|  | KEGG | 21,824 | 89.76 |
|  | Swissprot | 23,012 | 94.65 |
|  | TrEMBL | 22,970 | 94.47 |
| unannotated |  | 315 | 1.30 |

### Genome features of the African pike

CpG islands (CGIs), which are a significant group of CpG dinucleotide repeats in genome regions, are functionally important for genomic studies. The CGIs were identified across the genome using CpGIScan^32^. Ultimately, 24,297 CGIs were identified with a total length of 15.5 Mb. A range of genome features including gene density, repeat content, GC content, and GGI content were summarized and depicted in Fig. 2a. The CpG density was found positively correlated with GC content, gene density, and repeat content (Fig. 2b), following a similar pattern observed in other published fish and mammals genomes^33-35^.

**Fig. 2.**
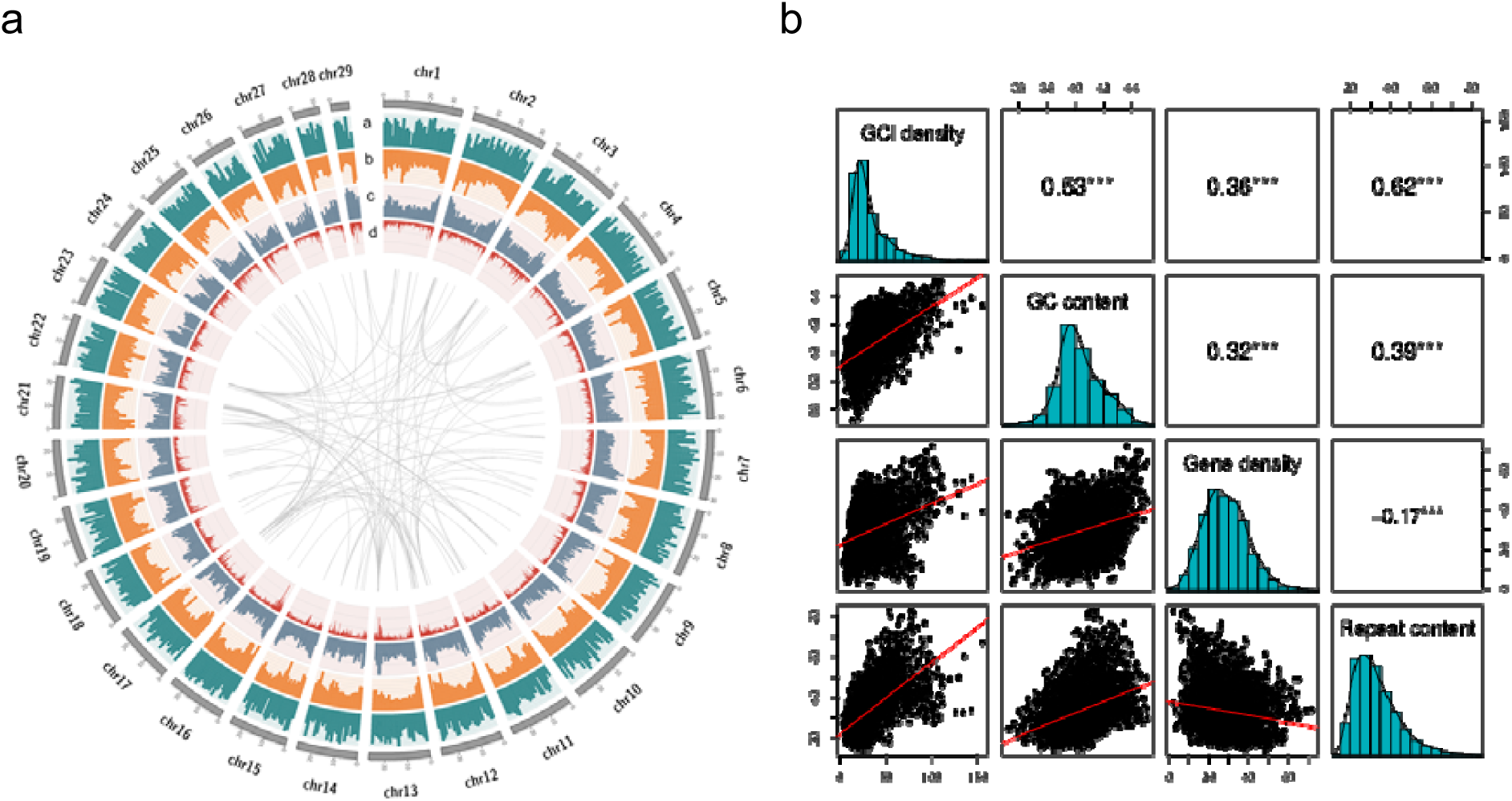
Genome features of the African pike. **(a)** The circos plot of 29 chromosomes in African pike. The tracks from outside to inside are: (1) Gene density, defined as gene counts per million base pairs. (2) Repeat content, defined as the proportion of repetitive elements within 1-Mb windows. It was presented in ratio as divided by the highest value. The axis ranges from 0 to 1. (3) GC content, quantified by the proportion of GC in unambiguous bases in 1-Mb window. It was presented in ratio as divided by the highest value. The axis ranges from 0 to 1. (4) CGI content, defined as CGI counts per million base pairs. The axis ranges from 0 to 200. **(b)** Correlation matrix among four genome features. The diagonal presents the distributions by histogram for corresponding genome features. The lower triangular matrix presents the bivariate scatter plots with a fitted linear model for each pair of genome features. The upper triangular matrix displays the Pearson correlation results plus significance level for the corresponding genome features. Different asterisks represent different significance levels: *p*-values 0.001 (***), 0.01 (**), 0.05 (*).

### Gene family identification

Gene family analysis among species provides significant insights into phylogenetic and evolutionary studies. The protein-coding genes from the African pike assembly, two sequenced species in order Characiformes (*A. mexicanus* and *P. nattereri*), and five other sequenced species including Atlantic salmon (Salmo salar), yellow catfish (*Tachysurus fulvidraco*), northern pike (*Esox lucius*), electric eel (*Electrophorus electricus*), and zebrafish (*Danio rerio*) were downloaded from NCBI database and analyzed. All-versus-all protein similarities were computed using BLASTP^26^ and the alignment results were used by TreeFam (v4.0)^36^ to deduce homologous gene sequences and identify gene families. Orthologue clustering analysis of predicted genes was conducted using MCL algorithm (Fig. 3a). Finally, we identified 9,661 gene families in the African pike genome (Additional file 1: Table S1). Compared to the two South American Characiformes (*A. mexicanus* and *P. nattereri*) and the zebrafish (*D. rerio*), 221 gene families were unique in African pike (Fig. 3b).

**Fig. 3.**
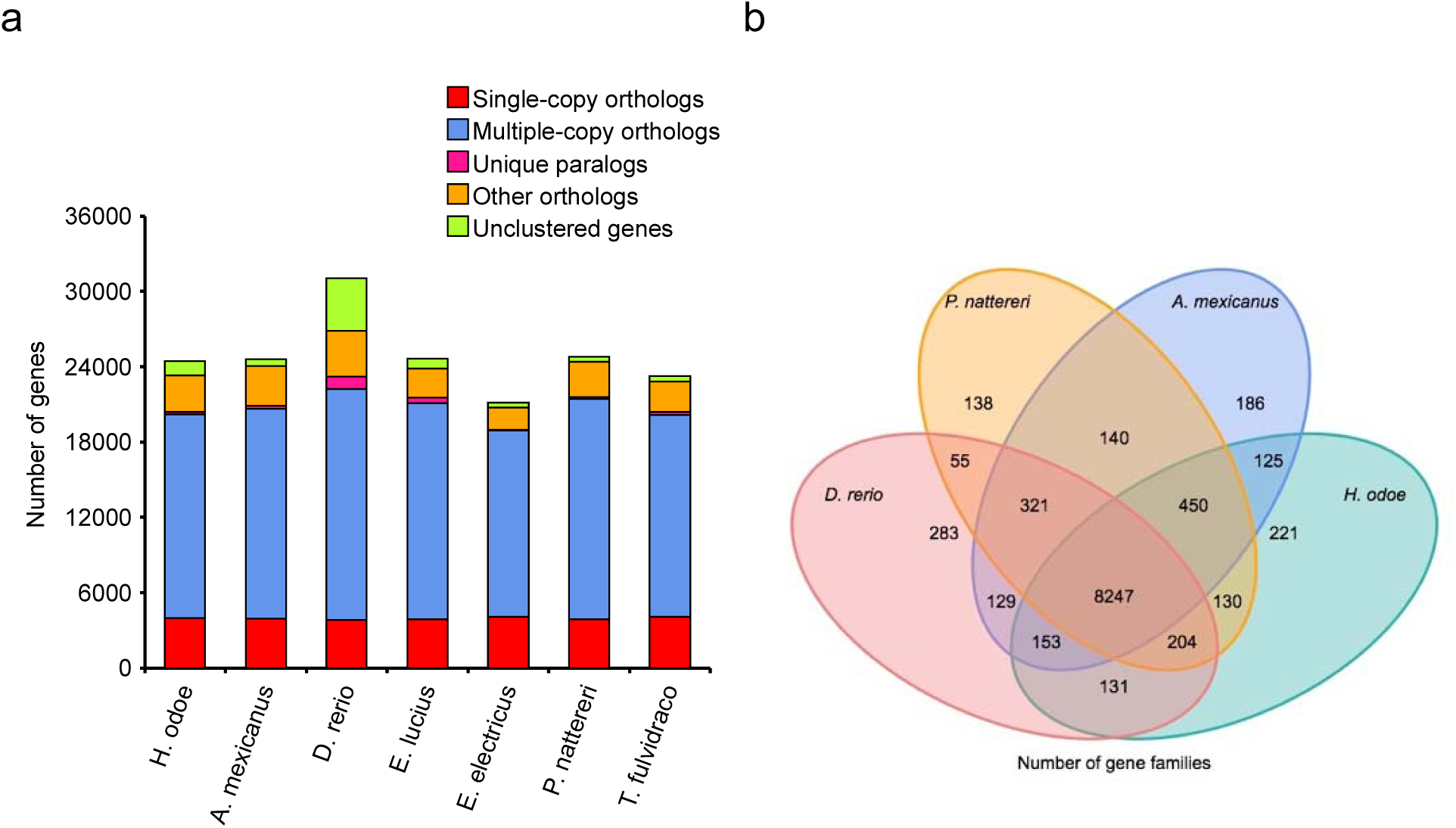
Comparative genome analysis. **(a)** Orthologue clustering analysis for African pike and other species. The x-axis displays the seven species and the y-axis presents the gene counts. Red refers to single-copy orthologs, blue refers to multiple-copy orthologs, pink refers to unique orthologs for corresponding species, orange stands for other orthologs, and green represents unclustered genes. **(b)** Venn diagram. Shared and unique gene families among the four species were shown in numbers in corresponding regions.

### Phylogenetic analysis

To study the evolutionary position of African pike, 3,106 single-copy genes from the above seven species were used for constructing phylogenetic tree and estimating divergence time. Protein sequences of single-copy gene families were aligned by MUSCLE (v3.8.31)^37^ and then were concatenated into a supergene matrix for each species. The alignment results were processed into PhyML (v 3.0)^37,38^ to construct a ML phylogenetic tree. Divergence time was inferred using the MCMCTree from the PAML package^39^. Divergence times from TimeTree database^40^ were applied for calibration, which include splits between *E. lucius* and *D. rerio* (198-211 Mya), between *D. rerio* and *T. fulvidraco* (170-183 Mya), and between *T. fulvidraco* and *E. electricus* (122-136 Mya). The phylogenetic tree showed that *H. odoe* was most closely related to *P. nattereri* with a divergence time around 73.8 Mya, and together the clade formed the sister group to *A. mexicanus* (Fig. 4a). The African family (*H. odoe*) was dispersed among the South American families (*P. nattereri* than *A. mexicanus*). Although sharing similar pike-like forms, the African pike was distantly related to the northern pike (*E. lucius*) with a divergence time around 205 Mya.

**Fig. 4.**
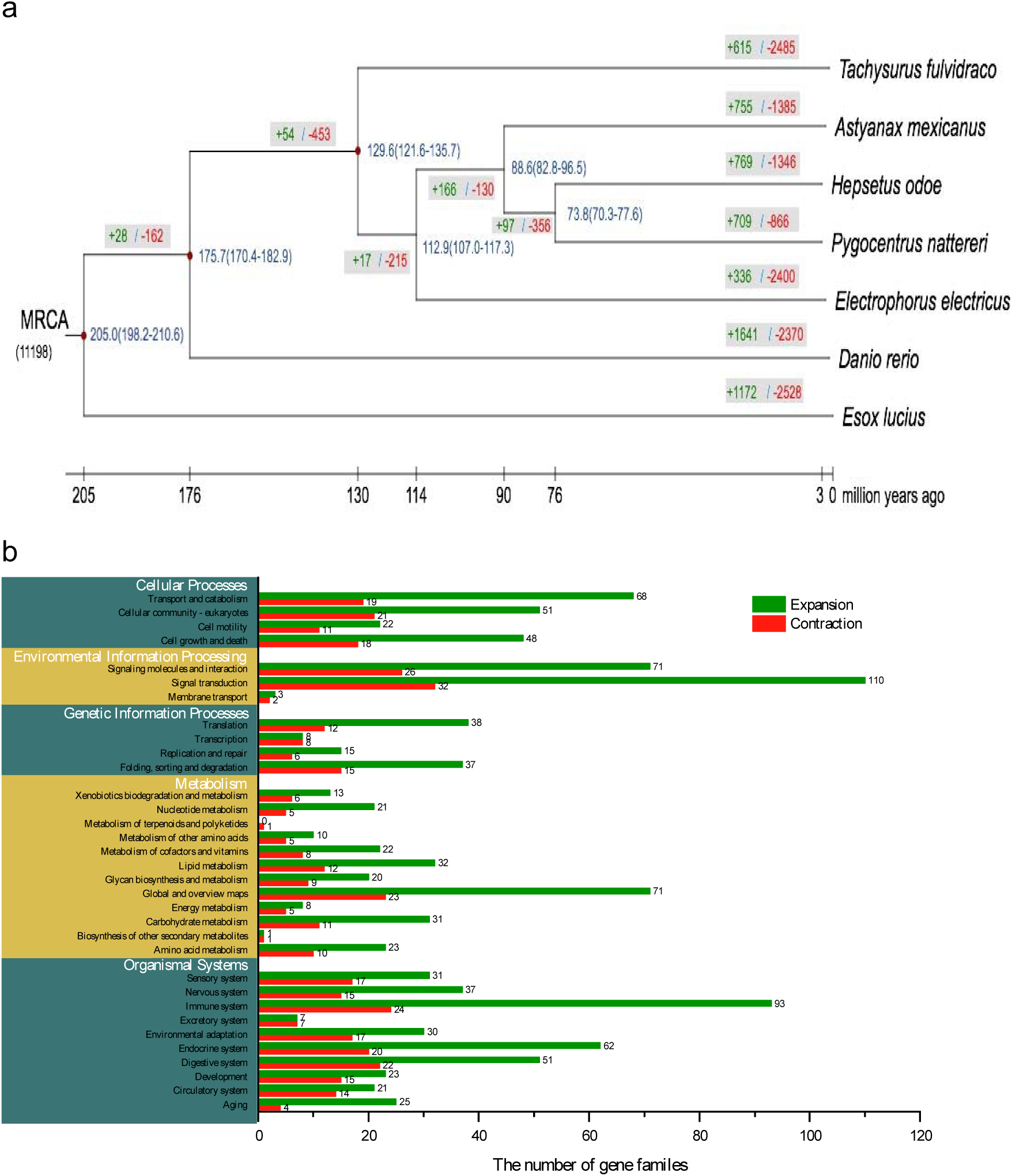
Phylogenetic tree, divergence time estimations, and the gene family expansions and contractions for African pike and other species from different fish orders. **(a)** The phylogenetic tree. Blue values represent the divergence time. Red nodes in the phylogenetic tree represent the reference divergence times. Green and red values represent expanded and contracted gene families for corresponding lineages, respectively. **(b)** KEGG functional enrichment of the significantly expanded and contracted gene families in the African pike.

### Expansion and contraction of gene families

Based on the gene family clustering results and divergence time estimation, we used Café (v2.1)^41^ to estimate the gene family expansion and contraction events during speciation. Results showed that in African pike genome 769 gene families were found expanded and 1,346 gene families were contracted (Fig. 4a). The 284 significantly expanded and 81 significantly contracted gene families (p< 0.05; Additional file 1: Table S2) in African pike were annotated with KEGG ortholog functions. Among that 93 (32.7%) gene families were strikingly expanded in immune system (Fig. 4b).

## Data Records

The sequencing data and genome assembly of the African pike were deposited in NCBI under BioProject accession PRJNA625402. The datasets reported in this study are also available in the CNGB Nucleotide Sequence Archive (CNSA: https://db.cngb.org/cnsa; accession number CNP0001012).

## Technical Validation

### Assessment of genome assembly

The contig and scaffold N50 of the African pike genome were 347.4 kb and 25.8 Mb respectively, with the longest scaffold 34,852,849 bp. We assessed the quality of the assembled genome using the Benchmarking Universal Single-copy Orthologs (BUSCO v3.0.2)^42^. The assembly reached 90.7%∼92.4% completeness compared to single-copy ortholog gene sets from atinopterygii, metazoans, and vertebrates in BUSCOs (Table 9). This demonstrates the high completeness of our genome assembly.

**Table 9.** Statistics of the BUSCO assessment.

| Types of BUSCOs | Gene Set |  |  | Assembly |  |  |
| --- | --- | --- | --- | --- | --- | --- |
|  | Number of<br>actinopterygii<br>(%) | Number of<br>metazoa<br>(%) | Number of<br>vertebrata<br>(%) | Number of<br>actinopterygii<br>(%) | Number of<br>metazoa<br>(%) | Number<br>of<br>vertebrata<br>(%) |
| Complete BUSCOs | 4,241<br>(92.5%) | 943<br>(96.4%) | 2,433<br>(94.1%) | 4,160<br>(90.7%) | 901<br>(92.1%) | 2,531<br>(90.6%) |
| Fragmented BUSCOs | 222 (4.8%) | 26 (2.7%) | 110 (4.3%) | 252 (5.5%) | 14 (1.4%) | 169<br>(6.5%) |
| Missing BUSCOs | 121 (2.7%) | 9 (0.9%) | 43 (1.6%) | 173 (3.8%) | 63 (6.5%) | 66 (2.6%) |
| Total BUSCO groups<br>searched | 4,584 (100%) | 978<br>(100%) | 2,586<br>(100%) | 4,584 (100%) | 978<br>(100%) | 2,586<br>(100%) |

Next, we evaluated the assembly of the twenty-nine chromosomes. The genome assembly was divided into 100kb bins. The signal for the interaction between any two bins was defined by the count of Hi-C reads covered by those bins, and the signal intensities were depicted in a heat map. The Hi-C heat map clearly split the bins into 29 blocks, and bins within the same chromosome showed substantially larger signal intensities than bins distributed on different chromosomes (Fig. 1). This demonstrates the high quality of the chromosome assembly.

### Gene prediction and annotation validation

Repetitive sequences in the assembly were masked before gene annotation. Gene model prediction in the African pike was realized by using a combination of *de novo* and homology-based approaches. Then the gene prediction results were integrated into a consensus gene set by GLEAN. Annotation completeness of the African pike gene set was assessed by BUSCO, reaching 92.5%∼96.4% completeness (Table 9). In addition, functional annotation of the predicted genes showed that 98.7% of them could be assigned into at least one functional term (Table 8). These results clearly indicate the annotated gene set is quite complete.

## Code Availability

All commands used in the analysis were executed by following the manual of the corresponding bioinformatics tools. There were no any custom specific codes.

## Acknowledgements

This work was supported by the special funding of “Blue granary” scientific and technological innovation of China (2018YFD0900301-05). We also thank for the technical supports from China National Genebank in stLFR library construction and sequencing.

## Author Contributions

G. F., H. Z., and X. L. designed the study. G. F., X. L., and W. Z. supervised the study. M. Z. and S. L. contributed to sample collection and sequencing experiments. X. D., X. Hong, S. S., X. Huang and B.O. performed bioinformatics analyses. X. D., X. Hong, and G. F. wrote the manuscript.

## Supplementary materials

### Additional file 1: Supplementary figures and tables

**Fig. S1.**
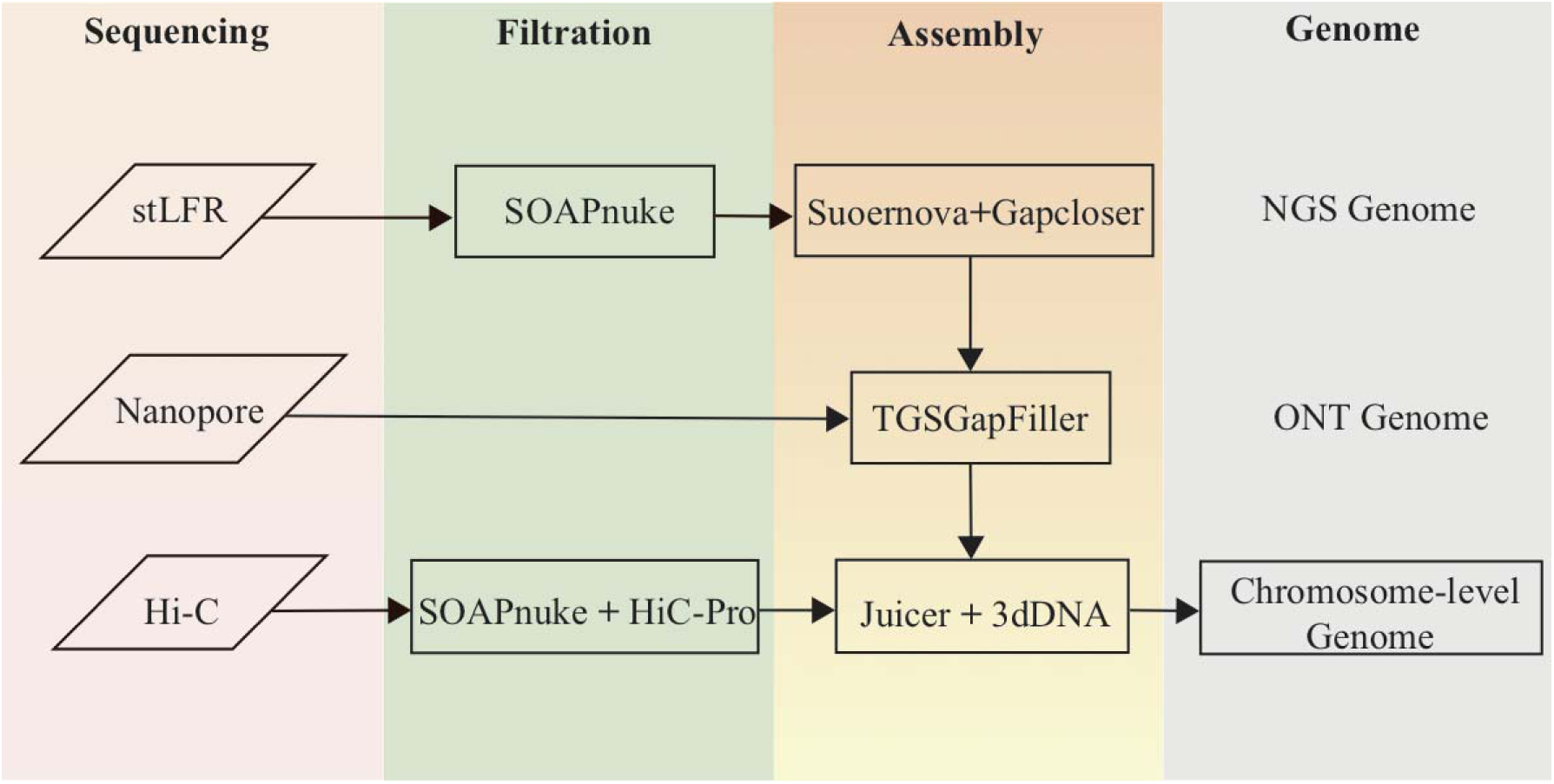
The assembly workflow.

**Fig. S2.**
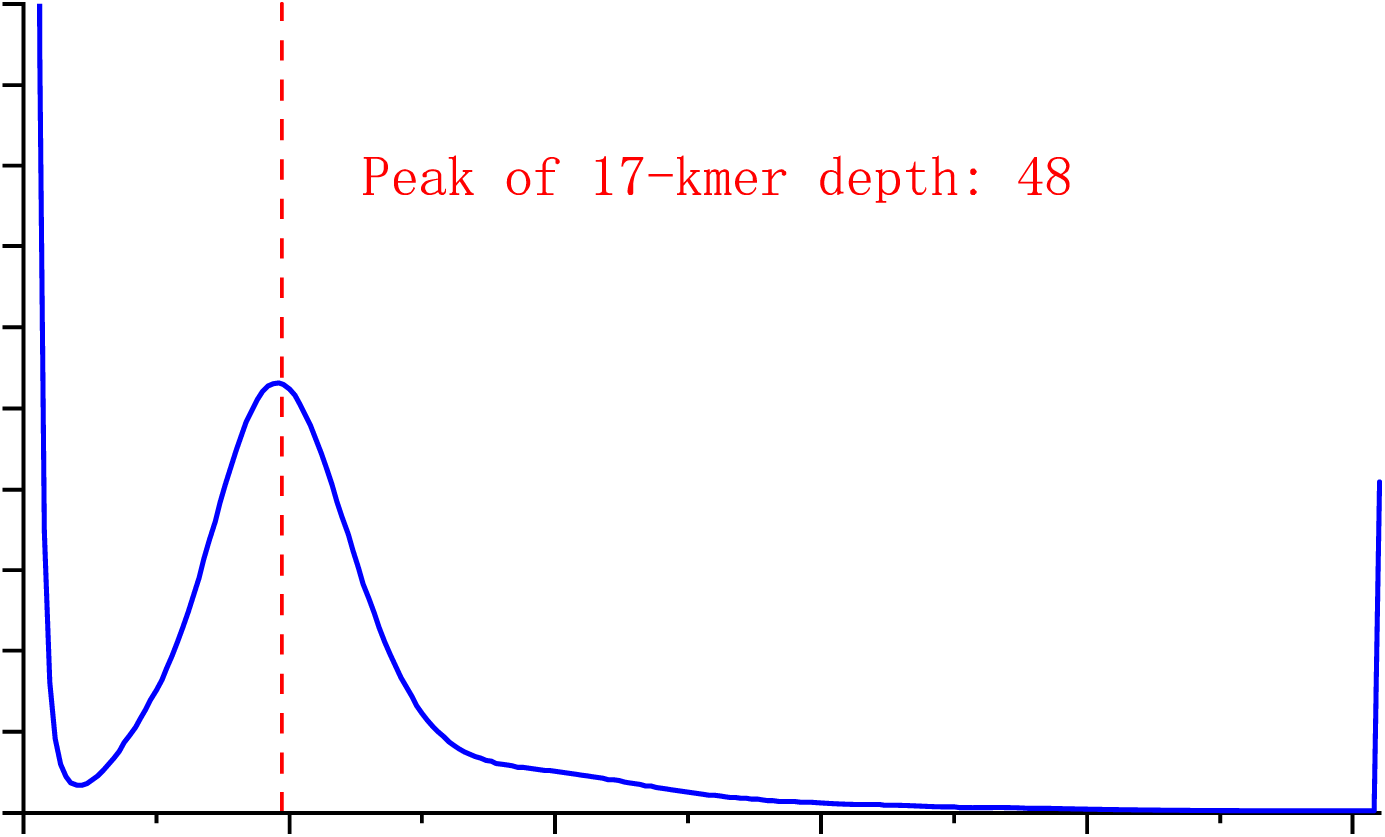
Distribution of k-mer frequency.

**Table S1.** Statistics of gene family clustering.

| Species | Genes number | Genes in families | Unclustered genes | Family number | Unique families | Average genes per family |
| --- | --- | --- | --- | --- | --- | --- |
| *H.odoe* | 24,314 | 23,300 | 1,014 | 9,661 | 103 | 2.47 |
| *A.mexicanus* | 24,612 | 24,043 | 569 | 9,751 | 73 | 2.82 |
| *D.rerio* | 31,056 | 26,849 | 4,207 | 9,523 | 155 | 2.82 |
| *E. lucius* | 24,657 | 23,851 | 806 | 9,358 | 119 | 2.55 |
| *E. electricus* | 21,162 | 20,761 | 401 | 9,243 | 18 | 2.25 |
| *P. nattereri* | 24,796 | 24,380 | 416 | 9,685 | 57 | 2.52 |
| *T. fulvidraco* | 23,258 | 22,796 | 462 | 9,389 | 78 | 2.43 |

**Table S2.**
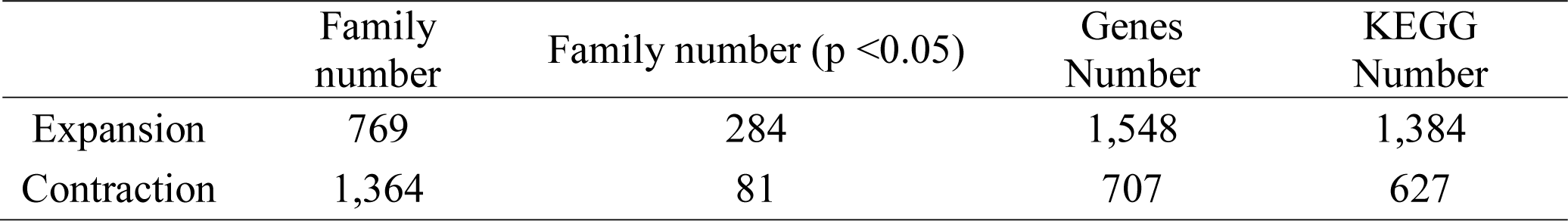
Gene family expansion and contraction statistics in African pike genome.

|  | Family number | Family number (p <0.05) | Genes Number | KEGG Number |
| --- | --- | --- | --- | --- |
| Expansion | 769 | 284 | 1,548 | 1,384 |
| Contraction | 1,364 | 81 | 707 | 627 |

